# SeroBA: rapid high-throughput serotyping of *Streptococcus pneumoniae* from whole genome sequence data

**DOI:** 10.1101/179465

**Authors:** Lennard Epping, Andries J. van Tonder, Rebecca A. Gladstone, The Global Pneumococcal Sequencing Consortium, Stephen D. Bentley, Andrew J. Page, Jacqueline A. Keane

## Abstract

*Streptococcus pneumoniae* is responsible for 240,000 - 460,000 deaths in children under 5 years of age each year. Accurate identification of pneumococcal serotypes is important for tracking the distribution and evolution of serotypes following the introduction of effective vaccines. Recent efforts have been made to infer serotypes directly from genomic data but current software approaches are limited and do not scale well. Here, we introduce a novel method, SeroBA, which uses a hybrid assembly and mapping approach. We compared SeroBA against real and simulated data and present results on the concordance and computational performance against a validation dataset, the robustness and scalability when analysing a large dataset, and the impact of varying the depth of coverage in the *cps* locus region on sequence-based serotyping. SeroBA can predict serotypes, by identifying the *cps* locus, directly from raw whole genome sequencing read data with 98% concordance using a *k*-mer based method, can process 10,000 samples in just over 1 day using a standard server and can call serotypes at a coverage as low as 10x. SeroBA is implemented in Python3 and is freely available under an open source GPLv3 license from: https://github.com/sanger-pathogens/seroba.

**DATA SUMMARY:**

1. The reference genome *Streptococcus pneumoniae* ATCC 700669 is available from National Center for Biotechnology Information (NCBI) with the accession number: FM211187
2. Simulated paired end reads for experiment 2 have been deposited in FigShare: https://doi.org/10.6084/m9.figshare.5086054.v1
3. Accession numbers for all other experiments are listed in Supplementary Table S1 and Supplementary Table S2.

**I/We confirm all supporting data, code and protocols have been provided within the article or through supplementary data files. ⊠**

**IMPACT STATEMENT:** This article describes SeroBA, a A-mer based method for predicting the serotypes of *Streptococcus pneumoniae* from Whole Genome Sequencing (WGS) data. SeroBA can identify 92 serotypes and 2 subtypes with constant memory usage and low computational costs. We showed that SeroBA is able to reliably predict serotypes at a depth of coverage as low as 10x and is scalable to large datasets.

## INTRODUCTION

*Streptococcus pneumoniae* (the pneumococcus) is a clinically important bacterium estimated to cause 700,000 to 1 million deaths in children under 5 years of age annually prior to the introduction of polysaccharide conjugate vaccines (O’Brien et al. 2009). The capsular polysaccharide biosynthesis (*cps*) locus, which encodes the serotype, is a major virulence factor in *S. pneumoniae.* The introduction of multi-valent pneumococcal conjugate vaccines has led to a substantial change in the circulating serotypes (Menezes et al. 2011) and decreased the number of deaths in children under 5 years of age to 240,000 - 460,000 annually (Wahl et al. 2016). By surveilling the circulating serotypes, the epidemiological trends of *S. pneumoniae* can be observed, pre- and post-vaccination. The rapid reduction in the cost of whole genome sequencing (WGS) has lead to its extensive use in the monitoring of pneumococcal serotypes (Lang et al. 2015)

To date there are nearly 100 known serotypes described for *S. pneumoniae* based on differing biochemical and antigenic properties of the capsule (Van Tonder et al. 2017). The *cps* locus, which encodes the serotype, can be very similar between serotypes from the same serogroup (such as serogroup 6) with some of them distinguished by a single nucleotide polymorphism (SNP), rendering a gene non-functional or altering the sugar linkage (Bentley et al. 2006). However, dissimilar loci may be grouped in the serogroup as they elicit a similar antibody response (e.g. serogroup 35). The large number of identified serotypes, and the high similarity between them, makes it challenging to computationally predict the serotype based on WGS data. Another challenge is recombination with other serotypes resulting in a mosaic *cps* locus (Salter et al. 2017) which may or may not affect the polysaccharide being produced. It is possible to have significant variation across the *cps* locus which does not lead to a different polysaccharide capsule being produced (Ko et al. 2013). Conversely novel serotypes can be generated through these processes and can go unnoticed by antibody-based serotyping (Geno et al.; Park et al. 2007). Finally, mixed populations in a single sample and contamination can lead to ambiguity.

There are a number of methods available to predict serotypes in *S. pneumoniae.* Besides the gold standard method, Quellung, which can be subjective in certain cases, there are five additional methods based on serological tests, at least eight semi-automated molecular tests based on PCR and one method that uses microarray data for serotyping (Jauneikaite et al. 2015). There are a number of insilico methods to detect the *cps* locus, which can then be used to predict serotypes from WGS data (Croucher et al. 2009; Leung et al. 2012; Kapatai et al. 2016; Metcalf et al. 2016). However, the tool described by Metcalf *et al.* is an in-house tool, and the tool described by Leung *et al.* only covers half of the known serotypes.

The only fully-featured automated pipeline for serotyping *S. pneumoniae* WGS data is PneumoCat, which was developed by Public Health England (PHE) (Kapatai et al. 2016). PneumoCat provides a capsular type variant (CTV) database including FASTA sequences for 92 serotypes and 2 subtypes as well as additional information about alleles, genes and SNPs for serotypes within specific serogroups. To predict a serotype, PneumoCat uses bowtie2 (Langmead and Salzberg 2012) to align reads to all serotype sequences. If the serotype belongs to a predefined serogroup or the serotype sequence could not be unambiguously identified, PneumoCat maps the reads to serogroup specific genes to identify the genetic variants. It is however computationally and memory intensive, and does not work with samples where there is a low depth of coverage in the *cps* locus region, as shown below.

To address these problems, we developed SeroBA, which makes efficient use of computational resources and can accurately detect the *cps* locus even at low coverage, and thus predict serotypes from WGS data using a database adapted from PneumoCAT (Kapatai et al. 2016). This accuracy was evaluated by comparing the results to a standard, validated dataset from PHE (Kapatai et al. 2016). We showed that it is scalable and robust by calculating the serotypes of 9,477 samples from the GPS project, an ongoing global pneumococcal sequencing project, on commodity hardware. Simulated read data, with varying coverage over a known reference sequence, was used to show the minimum depth of coverage required to call a serotype.

## THEORY AND IMPLEMENTATION

SeroBA takes Illumina paired-end reads in FASTQ format as input as shown in Figure 1. Precomputed databases are bundled with the application that describe the serotypes. The first of these is a *k*-mer counts database for every serotype sequence produced by KMC (v3.0.0) (Kokot et al. 2017), the second is an ARIBA (v 2.9.3) (Hunt et al. 2017) compatible database for every serotype, and the third is a capsular type variant (CTV) database, including FASTA sequences for 92 serotypes and 2 subtypes, as well as additional information about alleles, genes and SNPs for serotypes in specific serogroups. These databases were adapted from PneumoCAT (Kapatai et al. 2016). A *k*-mer analysis is performed on the input reads, and the intersection is found between these *k*-mers and the precomputed *k*-mer database of serotypes. The *k*-mer coverage of the input reads over the serotype sequences is normalised to the sequence length of the serotype sequence and the serotype with the highest normalised sequence coverage is selected. This step identifies the possible serotype or serogroup and ARIBA is used to confirm the presence of the selected serotype from the raw reads. If a serogroup is selected, the *cps* sequence produced by ARIBA and serotype specific genes are aligned with nucmer (Kurtz et al. 2004) to find specific variants, such as presence/absence of genes, SNPs, or gene truncations as defined in the CTV. The output of SeroBA includes the predicted serotype with detailed information that led to the prediction, as well as an assembly of the *cps* locus sequences.

**Figure 1:**
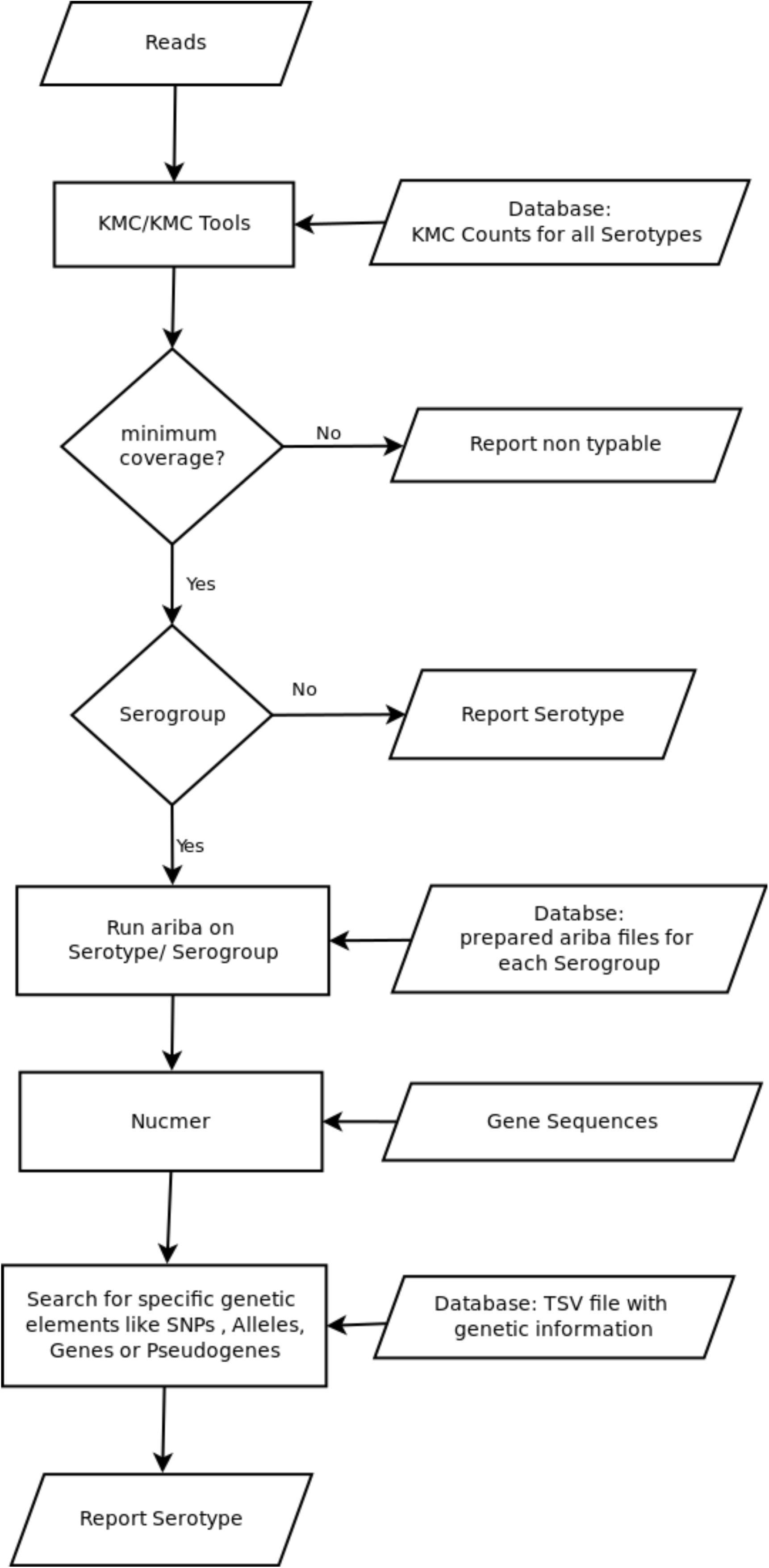
Flowchart outlining the main steps of the SeroBA algorithm

## VALIDATION DATASET

A validation dataset consisting of 2,065 UK isolates (Supplementary Table S1) retrieved from the PHE archive was originally used to evaluate PneumoCat. It consists of 72 out of 92 known serotypes, including all serotypes contained in commercial vaccines, and 19 non-typeable samples. The serotype of each sample was confirmed by latex agglutination with Statens Serum Institut typing sera (Kapatai et al. 2016). PneumoCat v1.1 (Kapatai et al. 2016) and SeroBA v0.1 were evaluated on an AMD Opteron 6272 server running Ubuntu 12.04.2 LTS, with 32 cores and 256GB of RAM. A single CPU (Central Processing Unit) was used for each experiment, repeated 10 times, with the mean memory usage and wall clock times noted.

Figure 2 summarises the serotypes called for each sample by each method. As serotyping with latex agglutination and Quellung can be subjective (Selva et al. 2012) and potentially imprecise, a serotype was said to be concordant if two or more methods agreed on the same serotype. This gave a concordance of 98.4% for SeroBA and 98.5% for PneumoCat with latex agglutination method. The reference sequences in the CTV for the serotypes 24A, 24B, 24F may not be representative for the circulating strains (Kapatai et al. 2016), so SeroBA will report serogroup 24 instead of reporting the serotype. As discussed in (Kapatai et al. 2016) serological prediction in serogoup 12 were error-prone, so a prediction of either 12B or 12F were counted as concordant.

**Figure 2:**
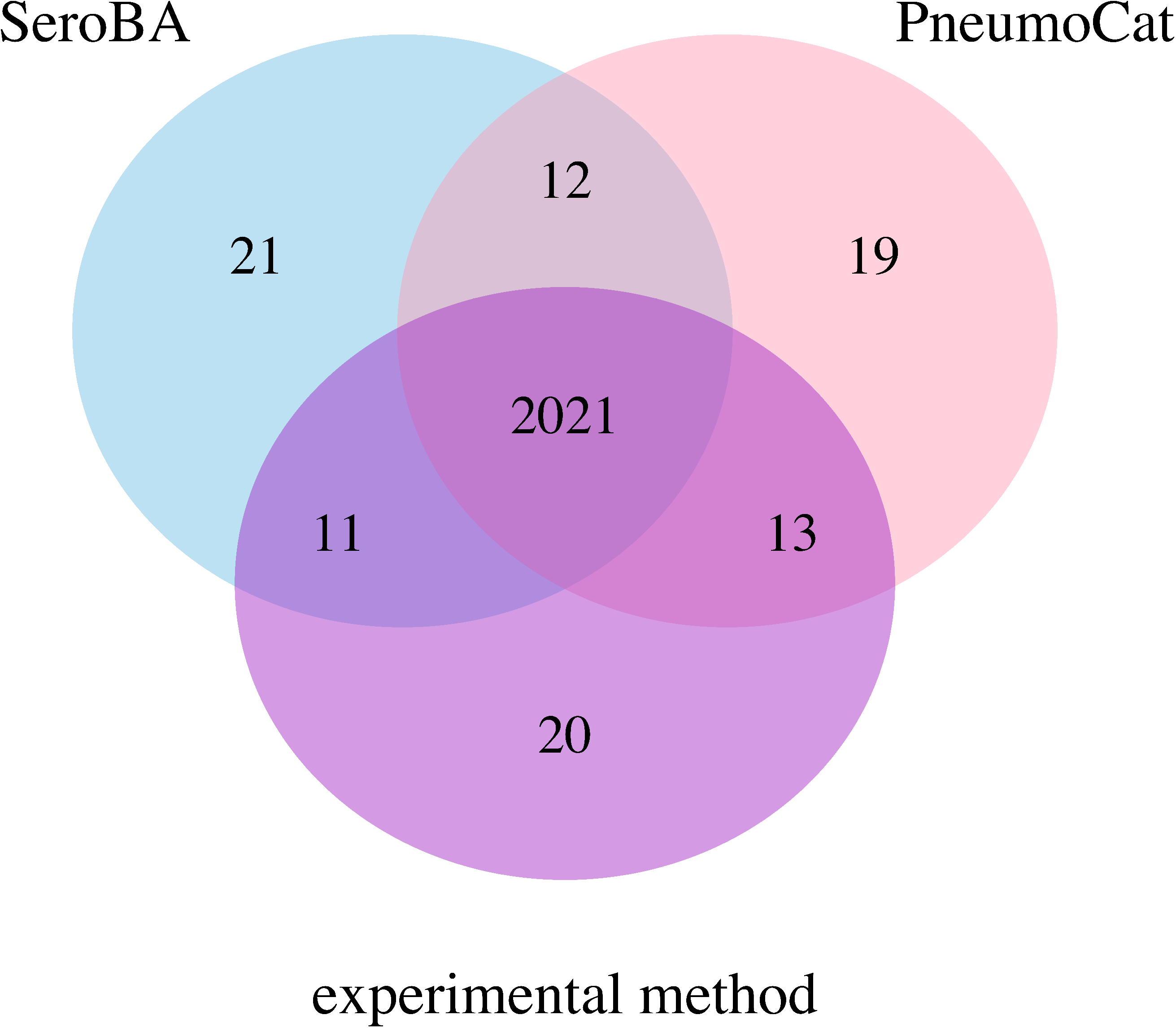
Agreement of serotyping results between different methods

The overall computational resources required to call the serotypes differed substantially between PneumoCat and SeroBA (Table 1): SeroBA was fifteen times faster and required five times less memory than PneumoCat.

**Table 1:** Performance of SeroBA and PneumoCat on the validation set

| Tool | Mean Wall Clock Time (m) | Mean RAM usage (MB) |
| --- | --- | --- |
| PneumoCat | 65.84 | 922.89 |
| SeroBA | 4.53 | 187.82 |

## EVALUATION USING A LARGE DATASET

To show the scalability of SeroBA to large datasets, we took 9,477 *S. pneumoniae* samples from the GPS project (Supplementary Table S2) and calculated the serotypes using the hardware setup previously described. A comparison with serotypes determined using experimental methods gave an accuracy of 98.2% for SeroBA. The serotypes were determined by different experimental methods as listed in Supplementary Table S2. Using all 32 cores resulted in a total wall-clock time of 823.78 hours. This showed that SeroBA can robustly scale to large datasets.

## IMPACT OF DEPTH OF COVERAGE

The effect of depth of coverage on the serotyping results produced by SeroBA and PneumoCat was evaluated by simulating perfect paired end reads over the serotype 23F *cps* locus from the *Streptococcus pneumoniae* ATCC 700669 (accession code: FM211187) reference genome (Croucher et al. 2009). Flanking regions of 1,000 bases were included on either side of the *cps* locus to eliminate confounding effects of low coverage at the locus boundaries. The reads with a length of 125 base pairs were generated by FASTAQ (v3.15.0) (https://github.com/sanger-pathogens/Fastaq) with an insert size of 500 bases and standard deviation of 50 with varying depth of coverage from 1x to 50x and from 100x to 350x in steps of 50. SeroBA started to predict serotype 23F at a depth of coverage of 10x while PneumoCat required nearly twice as much, needing at least 19x coverage. The computational resources required by SeroBA remained constant with increasing depth of coverage; however, the computational resource requirements of PneumoCat continue to grow linearly (Figure 3). At 350x coverage, PneumoCat took 3 times longer than SeroBA. Similarly, the amount of memory required by SeroBA stabilised at 150MB, regardless of coverage, whereas PneumoCat’s memory requirement grew with the depth of coverage, requiring 3 times more than SeroBA at 350x coverage. Each experiment was repeated 10 times and the mean was calculated.

**Figure 3:**
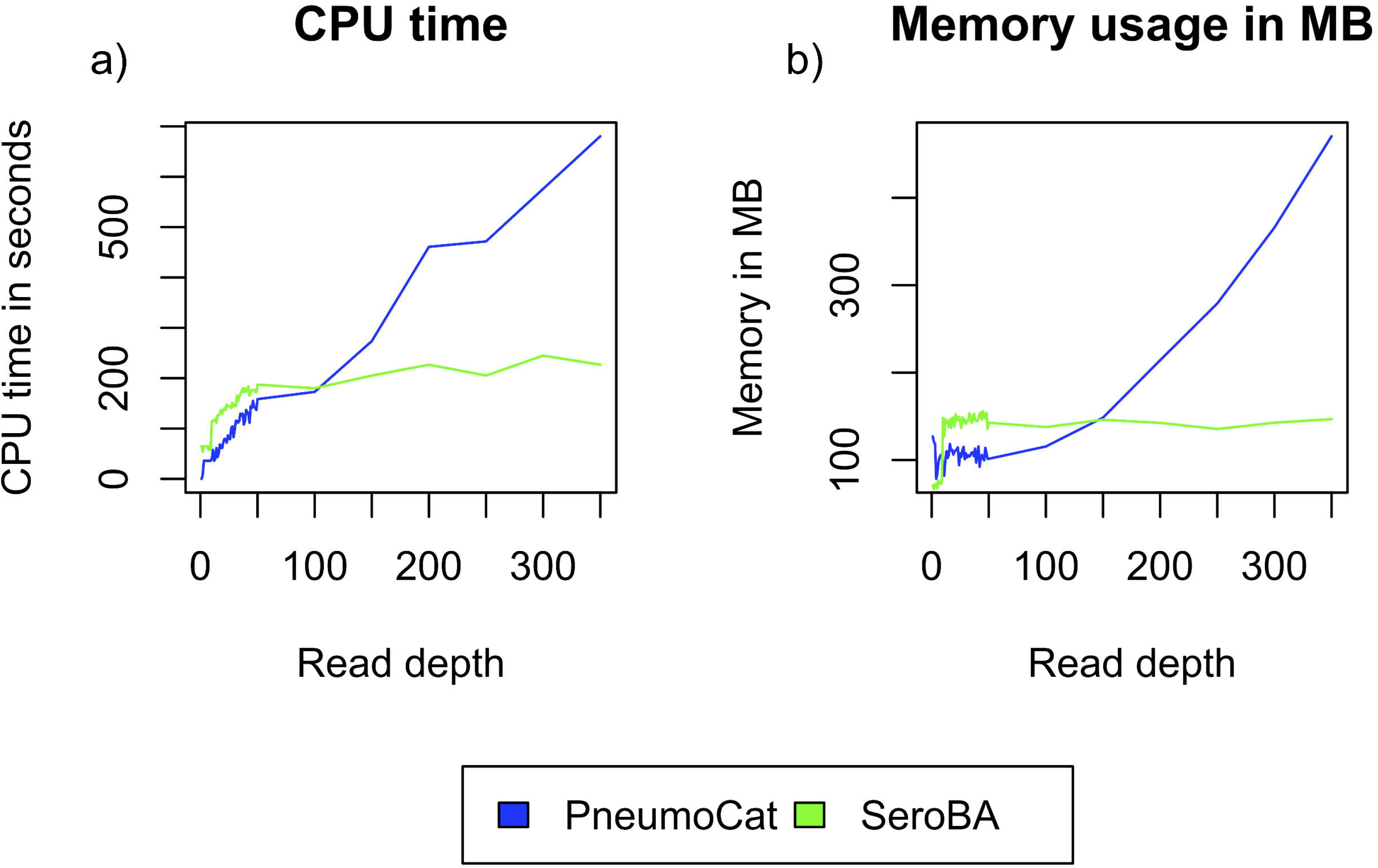
a) mean CPU time in seconds used by SeroBA and PneumoCat when varying the coverage from 1x to 350x; b) maximum memory allocation of SeroBA and PneumoCat when varying the coverage from 1x to 350x. Each data point represents the mean value of ten identical experiments.

## CONCLUSION

In this paper, we described SeroBA a method for predicting serotypes from *S. pneumoniae* Illumina NGS reads. We compared SeroBA and PneumoCat to a gold standard experimental serotyping method and showed that they had approximately the same level of concordance. However, SeroBA was fifteen times faster and required five times less memory than PneumoCat. The assembly of the *cps* locus sequence provides by SeroBA is another key feature that is very useful for further analyses and reference free comparisons. SeroBA was able to predict the serotype from only 10x read depth and scaled well on a large dataset of nearly 10,000 samples with a prediction accuracy of over 98%.

## AUTHOR STATEMENTS

[REMOVED FOR BLIND REVIEW]

## ABBREVIATIONS

SNP: Single nucleotide polymorphism
WGS: Whole genome sequencing
CTV: Capsular Type Variant database
CPS: Capsular polysaccharide biosynthesis
GPS: The Global Pneumococcal Sequencing

## DATA BIBLIOGRAPHY

1. https://github.com/sanger-pathogens/seroba
2. Lennard Epping, figshare. DOI: https://doi.org/10.6084/m9.figshare.5086054.v1
3. *Croucher, N. J, Streptococcus pneumoniae* ATCC 700669. NCBI., FM211187

